# A combined RAD-Seq and WGS approach reveals the genomic basis of yellow color variation in bumble bee *Bombus terrestris*

**DOI:** 10.1101/2020.08.12.248740

**Authors:** Sarthok Rasique Rahman, Jonathan Cnaani, Lisa N. Kinch, Nick V. Grishin, Heather M. Hines

## Abstract

**Background:** In the model bumble bee species *B. terrestris*, both males and females exhibit black coloration on the third thoracic and first metasomal segments. We discovered a fortuitous lab-generated mutant in which this typical black coloration is replaced by yellow. As this same color variant is found in several sister lineages to *B. terrestris* within the *Bombus s*.*s*. subgenus, this could be a result of ancestral allele sorting.

**Results:** Utilizing a combination of RAD-Seq and whole-genome re-sequencing approaches, we localized the color-generating variant to a single SNP in the protein-coding sequence of a homeobox transcription factor, *cut*. Sanger sequencing confirmed fixation of this SNP between wildtype and yellow mutants. Protein domain analysis revealed this SNP to generate an amino acid change (Ala38Pro) that modifies the conformation of coiled-coil structural elements which lie outside the characteristic DNA binding domains. We found all Hymenopterans including *B. terrestris* sister lineages possess the non-mutant allele, indicating different mechanism(s) are involved in the same black to yellow transition in nature.

**Conclusions:** *Cut* is a highly pleiotropic gene important for multiple facets of development, yet this mutation generated no noticeable external phenotypic effects outside of setal characteristics. Reproductive capacity was observed to be reduced, however, with queens being less likely to mate and produce female offspring, in a manner similar to workers. Our research implicates a novel developmental player in pigmentation, and potentially caste as well, thus contributing to a better understanding of the evolution of diversity in both of these processes.

## Background

Understanding the genetic architecture underlying phenotypic diversification has been a long-standing goal of evolutionary biology. In early research, discoveries and understanding of the genetic basis of traits relied on fortuitous mutant phenotypes predominantly in *Drosophila*. ([1], [2]) and subsequently in other model organisms (reviewed in [3]). These studies have not only contributed a myriad of insights about the characterization, chromosomal arrangement, and functional interactions of involved genes but have also ultimately shed light on the complex genomic mechanisms involved in natural variation [4]. Many of these early-era forward genetics studies [5] involved color-variable mutants. Such color variants have led the way in our understanding of evolutionary genetic processes, as coloration tends to be under strong selection and thus exhibits substantial and sometimes complex variation [6].

In recent years, the emergence of increasingly cheaper high-throughput sequencing techniques and availability of genomic resources and computational tools have expanded investigation of the genomic basis of color traits to a wide range of non-model organisms. Application of high-throughput sequencing (e.g. Whole Genome Sequencing, RNA-Seq, RAD-Seq) has provided solutions to many practical complications (e.g. knowledge gap in trait heritability, insufficient pedigree data, the infeasibility of lab-rearing or crossing) that previously limited the ability to unravel the genomic basis of color traits in many non-model organisms [7]. While some of this research has identified genes which have recurrent pigment-related roles in model organisms, many of these studies have provided novel insights about pigmentation and its key players (reviewed in [6], [8]). Such research has also contributed to broad principles in evolutionary genetics through revealing the multiple genomic routes (e.g., acting on *cis* and *trans*) to color variation (reviewed in [6], [9]; [10]) in both model and non-model organisms. These studies have also revealed the importance of co-option of major developmental genes for color patterning (e.g., [11], [12]), and the role of linked genetic variants in facilitating complex mimetic color phenotypes (e.g., [13], [14]).

Bumble bees exhibit an astounding diversity of color patterns, with the ∼260 species [15] of this genus displaying >400 color patterns ([16], [17]). This striking diversity has been largely attributed to the repeated divergence and convergence in color patterns onto numerous local Müllerian mimicry complexes, however, other ecological factors, such as thermoregulation and crypsis, may also be involved [16]. color is imparted in the thick setal pile (pubescence) on the head, thorax and abdominal sclerites in these bees, and is highly modular, with transitions between several colors (e.g. black, red, white, orange, yellow) across segments and many of the possible conceivable segmental combinations of these colors occurring across the lineage [17]. This phenotypically diverse genus is an emerging model system in evolutionary research [7], as this system contains ample polymorphisms that enable discovery of the genetic basis of coloration in natural populations ([18], [12]), can reveal evo-devo processes in segmental modularity [12] and can disentangle the microevolutionary processes involved in sorting allelic variation through its abundant natural replicates of identical segmental color transitions ([12], [7]).

*Bombus terrestris* is one of the most abundant bumble bee species of temperate regions of the western Palaearctic. As such, it is a major commercially reared pollinator utilized globally for greenhouse pollination services ([19], [20]), a development which has facilitated its use in laboratory studies. *B. terrestris* has become the leading model bumble bee for research in such areas as social evolution ([21], [22]), learning ([23], [24]), foraging behavior [25], immunology [26], invasive biology [27], ecological and landscape genetics [28], flight [29] and thermoregulation [30]. In recent years, research on *B. terrestris* has been facilitated by the availability of multifaceted genomic resources, as it has a publicly available annotated genome [31] and numerous additional available genomic [32], transcriptomic [33], and chromatin accessibility datasets [34].

*B. terrestris* belongs to the type subgenus *Bombus s*.*s*. [35], a lineage that has long been contentious regarding species status of many members of the complex. Part of this complexity has resulted from the color polymorphisms exhibited by the lineage, which have led to false inferences regarding species boundaries ([36], [37]). One of the most variable regions in coloration across the subgenus is the regional module involving the third thoracic (mesosomal) segment, first metasomal segment, and the posterior thoracic pleuron. This region transitions between black and yellow with intermediacy for some species, and can vary by species, region, and sex across this subgenus (Figure 1D). Examination of the color transition patterns of bumble bee species worldwide has identified the particular segments and colors that change most often in these bees ([16]; [17]). These studies found that yellow to black transitions are common, and that the posterior mesosomal and anterior metasomal segments are the segments most likely to vary in coloration across species ([16], [17]). *B. terrestris* also exhibits color variation, with nine regional color forms which are recognized as distinct subspecies ([38]; [39]), however, in all of these forms both males and females of this species have a characteristic black coloration in the abovementioned module (Figure 1). During rearing of inbred lines of *B. terrestris dalmatinus*, obtained from Israeli stock populations, a bee containing yellow in the aforementioned segments that typically contain black coloration was produced and subsequently inbred to develop a yellow mutant line. This fortuitous mutant enables assessment of the genetic basis of this trait using laboratory crosses.

**Figure 1:**
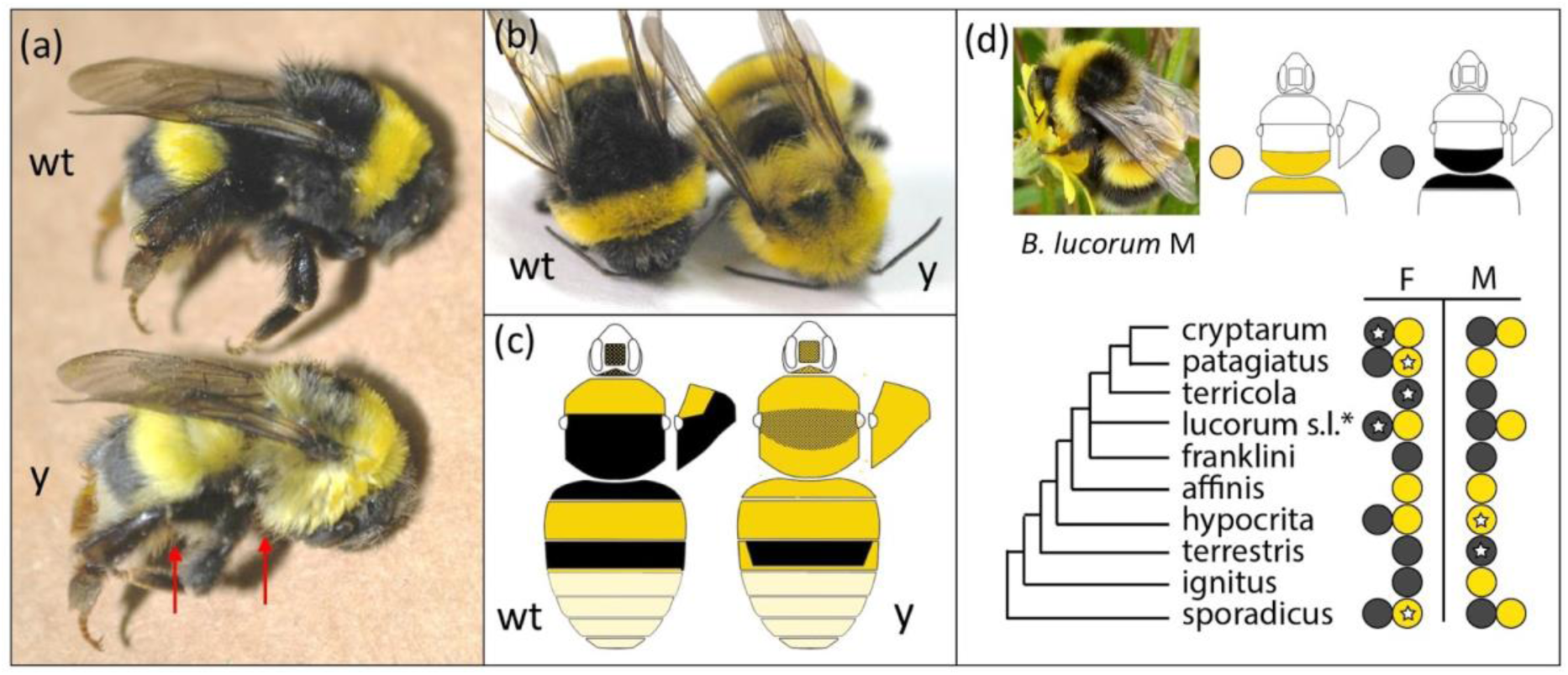
Wildtype and Yellow mutant color variants of *B*. *terrestris*. (a) Lateral view of wt and y workers. Red arrows point to yellow leg setae and an example of longer yellow setae in locations that have gained yellow color. (b) Dorsal view of workers. (c) Diagram of coloration pattern differences found in both morphs, showing the pattern for males, which matches the patterns for females. Stipules indicate mixed color. (d) The incidence of this phenotype (black versus yellow patterns on the third thoracic plus first metasomal segment) in the *Bombus s*.*s*. clade containing *B. terrestris*. Some species are polymorphic in this color and species vary in whether this color is sex-specific (male (M), female (F)). White stars in circles indicate specimens sampled for this study. Intermediates with mixing of setal colors in these segments can sometimes be found. *B. lucorum s*.*l*. Includes a complex of species, to which color assignment has not been ascribed [35]. The inset photograph (cropped from [115], creative commons license) depicts a yellow *B. lucorum* male morph from Scotland, which displays the same color pattern as the mutant *B. terrestris* phenotype. Phylogeny based on [116].

Recent studies ([40]; [41]) have identified black and ferruginous coloration in bumble bees to be imparted by melanin pigments (black eumelanin vs. ferruginous pheomelanin) [40], while yellow pigments likely involve both a pheomelanin [41] and a novel pterin-like pigment that occurs in yellow setae across bumble bee species [42]. Given the shift in melanin composition, candidate genes from the *Drosophila* melanin pathway [43] could be implicated in this color variation. A core set of key developmental transcription factors [e.g., *bric à brac (bab), wingless (wg), Distal-less (Dll), Abd-B, engrailed (en), doublesex (dsx)*] has been linked to segmental variation in *Drosophila* pigmentation [43], thus upstream developmental modulators could also be implicated in our case, especially given the segment-specific location of the phenotype. Tian et al. [12], for example, found that segment-specific variation involving red or black in mid-abdominal bumble bee segments was driven by cis-regulatory homeotic shifts in the posterior segmental *Hox* gene *Abd-B*. In *Drosophila*, morphological characteristics of the abovementioned set of segments are determined by the *Hox* gene, Ultrabithorax *(Ubx)* [44]. The localization of *Ubx* could explain why this region operates as a module. There are many additional developmental players that have been implicated in spatially preprogramming color pattern elements in butterflies [6] and which could hypothetically program segmental patterning in these bees [45]. Unravelling the SNPs/genes driving this mutant yellow coloration can expand our understanding of genetic targets for driving pigmentation and segmental differences, and potentially reveal underlying genetic processes by which the inheritance of this black to yellow shift occurred within the entire *Bombus s*.*s*. and broader bumble bee lineage.

In our current research study, we aim to unravel the genomic basis of mutant yellow coloration in *B. terrestris*. To achieve this goal, we utilize a combination of genome-wide reduced-representation approach (RAD-Seq) and whole-genome resequencing data on crossed offspring of both color phenotypes, which we analyze using genotype-phenotype association analyses to identify the genomic region associated with the black to yellow transition. We then investigate the potential function of the implicated mutation by comparing the predicted protein structure elements between wildtype and mutant protein sequences. Finally, we determine the evolutionary history and degree of sequence conservation of the identified genomic region in closely related species and across all Hymenopterans to better understand its potential role beyond this lab-generated *B. terrestris* mutation. Utilizing an inbred lab mutant line in a manner similar to early-era forward genetic approaches in model systems and harnessing the power of next-generation genome sequencing and growing genomic resources in non-model organisms, this study reveals a key developmental gene involved in pigmentation pathways in arthropods which could serve as a candidate gene for further research on extensive color pattern diversity in bumble bees.

## Results

### Phenotypic assessment

The mutant yellow and wildtype black color forms included in this study are outlined in Figure 1a-c. Although changes in color are most notable in the third mesosomal segment and first metasomal segment, mutant forms have increased in yellow hairs in several body regions, which in some cases is accompanied by longer and more dense setae. In the head there is an approximately 50% mix of yellow plumose (longer branches) setae mixed with shorter-branched black setae in the face as well as posterior to the eyes in the mutant form, and the setae in these regions appear longer and more dense. The wildtype bees have nearly all black hairs in these regions with fewer plumose setae and no yellow setae behind the compound eyes. The top of the head (vertex) is mostly yellow in the mutant form and mostly black in the wildtype. In the thorax (mesosoma), the mutant form has yellow in the third thoracic segment, yellow and black setae mixed in the second segment, yellow on the pleuron, and yellow in the ventral region. These setae appear longer than those in the wildtype. The wildtype instead is fully black in the second and third segments and nearly all black in the ventral thorax and the thoracic pleuron. In the metasoma, the first segment is yellow in the mutant and black in the wildtype. The third segment is mostly black with a thin yellow line at the distal boundary in the wildtype but this distal yellow line is thicker and runs up the lateral parts of the segment in the mutant form. The venter (sternites) has more and thicker and longer yellow hairs in the mutant form. The mutant form also has longer, thicker, and more yellow hairs on the femur and parts of the tibia, whereas the wildtype has only black hairs in these parts. Aside from the head, where there are more plumose hairs in the mutant form, the composition of setal types (short vs. long branched setae) is similar in both forms in other parts of the body. No morphological effects outside of setal traits were observed.

The yellow trait was deemed to be recessive, as heterozygous females are wildtype and her male offspring are 50% yellow form and 50% wildtype. While the initial mutant contained mostly pure yellow coloration in the respective segments, subsequent crosses reinforced the phenotype, making the yellow in the respective segments more pure (less intermediate/admixed with black). Accompanying this yellow mutation was a shift in female behavior. Yellow females showed reduced interest in mating with males of any phenotype or line, and low overall mating frequency (∼20% compared to ∼80% in wildtype).

### Gene localization: RAD-Seq and GWAS

Utilizing a combination of RAD-Seq and WGS approach on F2 offspring from crossed wildtype and mutant color forms, we narrowed the allele involved in the mutant yellow phenotype to a single nucleotide. First, 57,712 SNPs identified from the 90 RAD-Seq samples, revealed a single broad genomic region (∼6 Mb) on chromosomal scaffold NC_015770.1 of *B. terrestris* (Bter_1.0, GCA_000214255.1; [31]) which is highly associated with this color trait change (black vs. yellow) (Figure 2 a & b). As the RAD-Seq approach provides a reduced representation of genome-wide variation, it was not surprising that it failed to yield any fixed SNPs, thus we utilized a whole-genome sequencing approach (Fig 2 c & d) on 7 individuals of each phenotype that represent unique haplotypes from RAD-Seq data to target SNPs in the implicated interval from RAD-seq. GWAS within this narrowed scaffold (NC_015770.1) revealed a single fixed point mutation (C (mutant), G (wildtype); the genomic position 3802598) which falls at the top of the association peak from the RAD-Seq data. This SNP was determined to fall in the exonic sequences of the homeobox transcription factor *cut*. Our genotyping of this color locus across individuals utilized in the RAD-Seq analysis revealed a complete fixation of this SNP between wildtype and yellow mutants (data available in Dryad [to be provided upon publication]).

**Figure 2:**
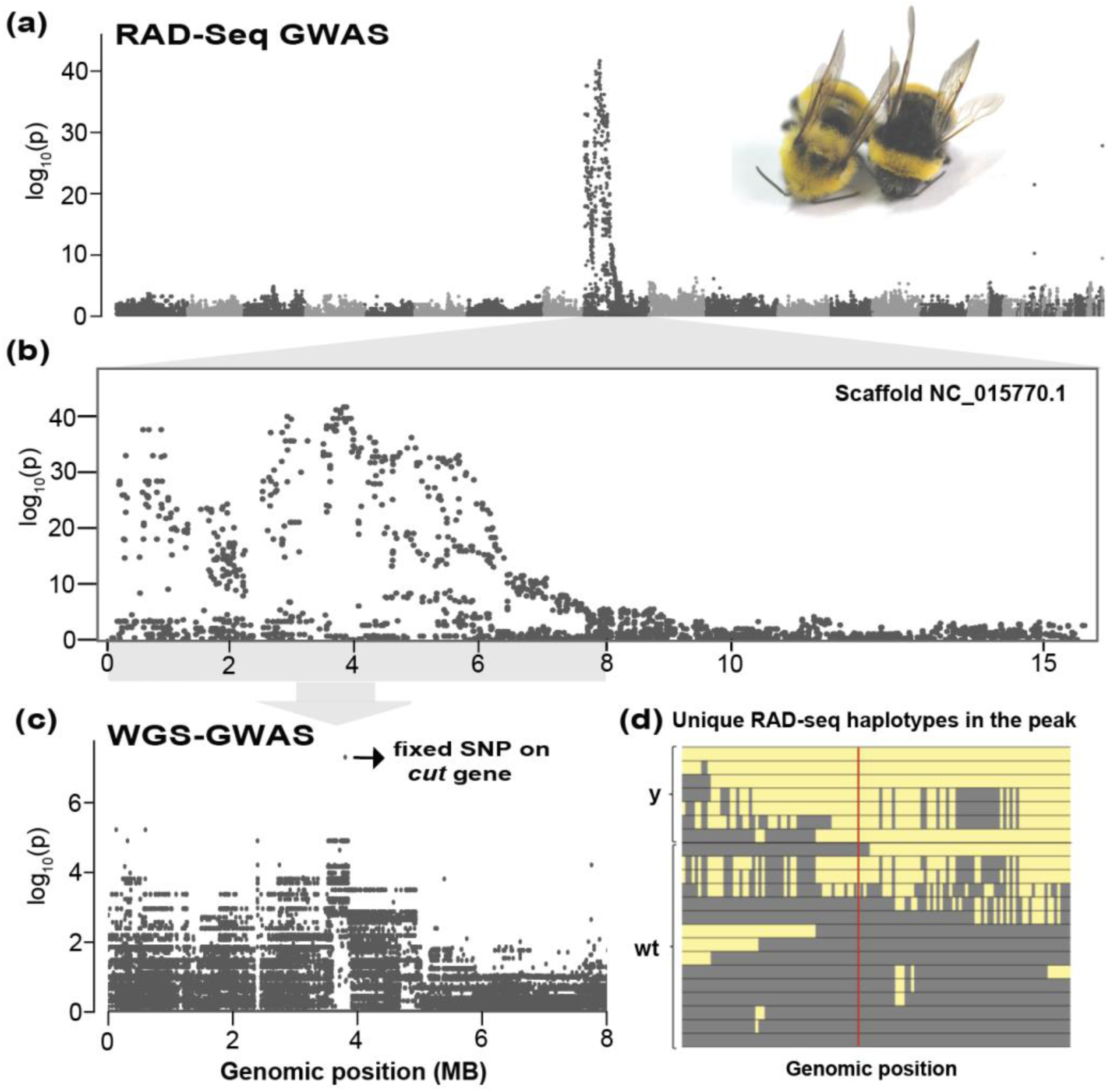
Identifying the color locus through a combination of WGS and RAD-Seq approach. (a) A genome-wide Manhattan plot using RAD-Seq data reveals a single broad peak of association in scaffold NC_015770.1. Scaffolds represented in different grayscale shades. (b) Zoomed view of the Manhattan plot for scaffold NC_015770.1 (c) A Manhattan plot using whole-genome sequencing data of the peak region from RAD-Seq. A single fixed SNP on *cut* gene is highlighted, (d) A diagram of the unique haplotypes observed across RAD-Seq data in the ∼3 MB (Genomic locations 2015753-5005430 on scaffold NC_015770.1) region of association used to select seven genetically distinct individuals for genomic sequencing of each color form. This is based on actual haplotypes of sampled individuals, focusing on a subset of SNPs that varied primarily between yellow mutant and wildtype individuals. SNPs are colored the color of the variant with the highest frequency of each allele (Yellow [y] for the mutant and gray for the wildtype [wt]). The red line indicates the position in the GWAS of the actual fixed SNP (not sequenced in RAD-Seq data). The filtered SNP dataset is deposited in Dryad digital repository [to be provided upon publication].

### SNP conservation and origin

All comparisons to other taxa outside of our samples revealed that they possess the wildtype allele in this position, thus supporting that this mutant is lab generated. Analysis of a publicly available whole-genome resequencing dataset of wildtype populations of *B. terrestris* [32] revealed only the variant present in our wildtype individuals (n=22; SNP dataset provided in Dryad digital repository [to be provided upon publication]). Genotyping of *cut* DNA sequences of multiple (n=7) members of closely related species belonging to the *Bombus s*.*s*. subgenus (Supplementary Table 1) revealed that all of these species also possess the “wildtype” allele, regardless of their yellow and black phenotypes (GenBank accession numbers: [to be provided upon publication]). A BLAST (blastn) search across all Hymenopterans (NCBI:txid7399; n=143) and analysis of aligned *cut* protein orthologs (n=40) obtained from OrthoDB database [46] (protein alignment file is provided in Dryad digital repository [to be provided upon publication]) also revealed no SNP or amino acid variation respectively, as all of them have the *B. terrestris* wildtype allele.

### Exploring the function of the implicated mutation

There are nine exons of the *cut* gene (total CDS length, 4806 bp) based on the predicted annotation (NCBI mRNA Reference Sequence: XM_012311834.2). The identified SNP [Wildtype(G) vs. Yellow mutant (C)] is on the fourth nucleotide of Exon 2 of *B. terrestris cut* mRNA, and induces a non-synonymous mutation [Wildtype (Alanine); Yellow mutant (Proline)] on the 38th codon position in the 1601 aa long protein (NCBI Reference Sequence XP_012167224.1). Our sequencing of *cut* cDNA (GenBank accession numbers: [to be provided upon publication], alignment available on Dryad [to be provided upon publication]) from wildtype and yellow mutant adult samples revealed that they both contained the SNP in their RNA, confirming they are part of the protein as expected based on automated annotations. This gene is not known to have natural transcript splice variants in *B. terrestris* according to its latest annotation, however, in other *Bombus* species present in NCBI and other members of Hymenoptera and Arthropoda, multiple splice variants, including at this boundary, are inferred. Amplified transcripts did not have variation in splicing and did not show signs of residual peaks in chromatograms at the splice boundary that would suggest partial production of alternative transcripts, however, as PCR could miss large insertions and rare products, more thorough transcriptomic analysis would be needed to test whether splice variants are involved.

### Protein Structure and conserved domain structure and the role of mutation

Figure 3a highlights the domain composition of the *cut* protein sequence, which includes two confidently predicted coiled coils (residues 30-68 and 90-148), followed by three *cut* domains (residues 526-598, 936-1008 and 1170-1239) and a homeobox domain (residues 1294-1347). The Ala38Pro variant maps to the N-terminal coiled coil. The Ala residue present in the wildtype sequence has a helical propensity consistent with the predicted coiled coil secondary structure (Figure 3a-b). However, mutation of this residue to Pro places a known helix breaking residue in the middle of the coil ([47]; [48]). As a result of this sequence change, the coiled coil prediction becomes less confident (Figure 3a and 3c). The helix breaking propensity of the Pro mutation results from an inability of the residue to complete the backbone hydrogen bonding pattern of an a-helix (Figure 3d-e). As such, coiled-coils infrequently contain Pro residues [49], and the introduction of a Pro residue into a coiled-coil can lower its helical content and disrupt its oligomeric state by introducing a kink into the helix [50].

**Figure 3:**
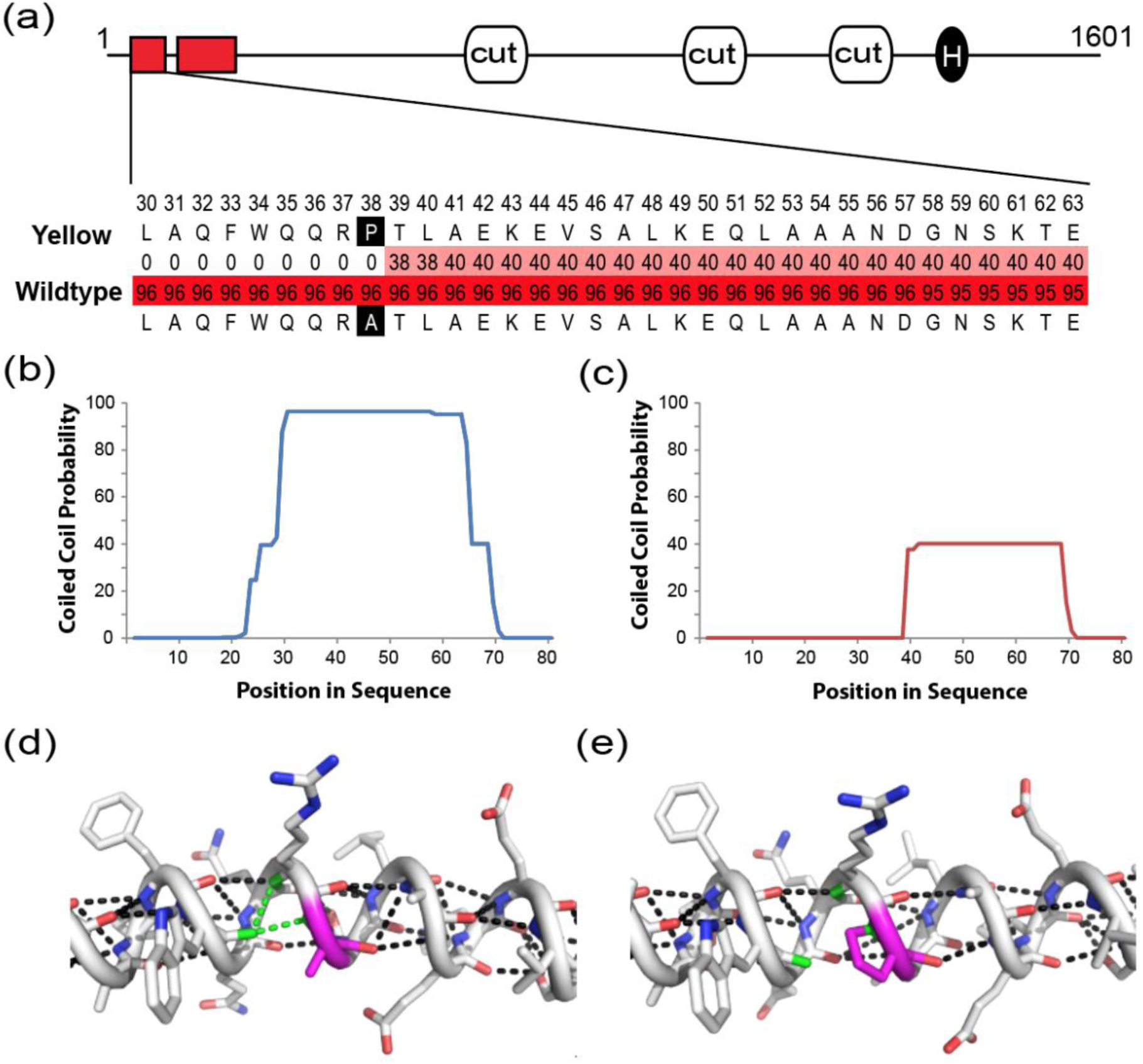
Predicting the structural differences between wildtype and yellow mutant *cut* proteins. (a) The *cut* protein sequence includes 3 *cut* domains (grey boxes) and a homeobox domain (light grey circle). Coiled-coil regions (MARCOIL probability >90) are indicated by red squares. MARCOIL probability list per residue for the wildtype (bottom) and yellow mutant (top) sequence (residues 30-63, labelled above) are highlighted from white to red in color scale from 0 to 100. MARCOIL probability plots for the N-terminal sequence (residues 1-80) of (b) wildtype and (c) Yellow mutant variant. (d/e) The *cut* coiled-coil (predicted from residues 20-55) is depicted in cartoon tube from the N-terminus (left) to the C-terminus (right), with residues in stick and colored according to atom type: carbon (white), nitrogen (blue) and oxygen (red); the wildtype sequence (d) can form helix stabilizing backbone hydrogen bonds (black dots) surrounding Ala38. Hydrogen bonds form between the Ala38 backbone nitrogen (and the i-1 Arg residue backbone nitrogen) and the i-4 Trp residue backbone carbonyl (green dots). The Ala38Pro mutation removes the ability to form a hydrogen bond between the backbone nitrogen and the i-4 carbonyl (colored green) in the yellow mutant variant (e) which destabilizes the helical propensity and can kink the coil.

Coiled coils often interact with each other to mediate functional protein interactions. Using physical interactions reported for *D. melanogaster* proteins in BIOGRID [51], we identified five high throughput interactions (*beag, nelf-A, nelf-E, brd8*, and *Bx42*) with *D. melanogaster cut* that localize to the nucleus and contain predicted coiled-coil sequence regions that could interact with the coil in *cut* (Supplementary Table 2).

## Discussion

Utilizing a combination of reduced-representation and whole-genome sequencing, we identified the color-controlling locus in the yellow mutant phenotype in *B. terrestris* to a single non-synonymous SNP in a homeobox transcription factor, *cut* (*ct*). *Cut* is a major developmental player that plays diverse roles in many different cells and tissue types including the brain, sensory organs ([52], [53]), wing discs, muscle tissues, Malpighian tubules, and reproductive organs [54], performing as a major selector gene of cell type and fate ([55]; [56]). The current study adds *cut* to a list of genes that can be involved in color patterning, and which may play a role in the diversification of color patterns in natural bumble bee populations.

Often, discovery of genes driving coloration identify pigmentation pathway genes (e.g., [57], [58], [59]). In this case, targeting the genetic basis of a color mutant has led to the discovery of a novel upstream player that can drive aspects of color patterning. It is hypothesized that color patterning along the body is likely dictated by regionally restricted developmental genes that can then be co-opted to drive spatial differences in pigmentation. For example, complex wing-spot color formation in *Drosophila guttifera* is driven by the co-option of a major developmental gene wingless which initially evolved to turn on pigmentation in the veins where it was expressed [60]. Using the pre-existing developmental pattern genes to generate novel and localized color-related phenotypes is a recurrent common theme in Lepidoptera wing coloration ([61], [62], [63], [11], [64], [65]). Several regionally restricted developmental genes (e.g.; *en, Omb, hh, ptc, wg*) have been hypothesized to play a major role in generating segment-specific abdominal pigmentation as well [65].

No clear link between *cut* and body-color pigmentation has been made previously, however, in *Drosophila*, analysis of a *cut* mutant (Dmel\ct^9b2^: FlyBase ID: FBal0028091; first discovered by Hannah [66] and later catalogued in [67]) found that this mutant lead to defective body color, with the fly exhibiting yellowish-tan pigmentation throughout the body. *Cut* may play a role in color patterning through its known interactions with other important developmental regional selector genes, such as *wingless (wg)* and *Notch (N)* [68]. In Lepidoptera, *cut* displays a high-level of co-expression with *wingless* in a spatiotemporal manner, acting as a “molecular cookie-cutter” to determine the complex wing shapes [69]. This *cut*/*wg* boundary determination mechanism in Lepidoptera is evolutionarily derived as it is different from the mechanisms used in *Drosophila* and other holometabolous arthropods and it may have facilitated the evolution of the astounding wing shape diversity in Lepidoptera [69].

A long-standing tenet in evolutionary developmental genetics is whether the evolutionary forces are more likely to act on cis-regulatory modules (CRMs) or trans-regulatory (protein-coding) factors. The classic school of thought in evolutionary developmental genetics has emphasized the pre-eminence of cis-regulatory element mutation ([70], [71]) in generating novel morphological phenotypes. Under this line of thinking, a protein-coding mutation at one of the most important developmental transcription factors should have widespread effects across multiple organs and systems as it is likely to have “deleterious” consequences. Many studies investigating the genetic variation of coloration have revealed the frequent targeting of cis-regulatory elements in highly pleiotropic genes (e.g., [72]; [43]) and protein-coding mutations in non-pleiotropic pigmentation genes (e.g., [73], [57]; [58]; [59]). The fitness consequences of *cut* protein mutation could potentially be substantial, as in *Drosophila* many documented *cut* mutants have displayed lethal or semi-lethal effects [74]. So, how does an apparently deleterious mutation on the protein-coding region of a highly pleiotropic gene such as *cut* have such limited phenotypic effects? In this case this novel mutation (Ala38Pro) occurs outside the characterized homeobox and *cut* domains. It affects protein structure instead through loss of a coiled-coiled structure near its N-terminus (Figure 3a). Coiled-coiled structures have been speculated to play important roles in protein-protein interactions ([75], [76]). For example, in *cux1* (cut-like homeobox1) proteins (a gene family in the *cut* superclass and includes *cut* genes in Arthropods) the gain of a coiled-coiled structure in N-terminal region by alternative splicing ([75]) results in alternative localization of the protein to Golgi bodies where they act as a transport protein [77]. In the present case, it can be assumed that this mutation has little or no impact on essential developmental functions of the genes and likely does not alter all of its roles, as it is not lethal for *B. terrestris* mutant types and has limited phenotypic effect. This leads one to contemplate whether *trans* effects necessarily exhibit high levels of pleiotropy, as altering specific domains of a protein may only affect some of the functions of a protein, for example those where specific proteins are present to interact with, thus generating the more localized tissue-specific effects that typically characterize cis-regulatory modifications. Indeed, there is a growing body of evidence in recent years demonstrating the importance of protein-coding mutations, which can act in a similar fashion as cis-regulatory mutations to generate modularity in gene regulation given that not all domains are functional in all contexts ([78]; [79]). Another possibility is that the localized effect of the *cut* gene is influenced by the levels of *cut* expression in implicated segments. For example, if the *cut* gene expression in that specific segment is close to the threshold levels needed to invoke specific responses or interaction, modifying a protein structure outside its characteristic protein domains can reduce its functionality and could result in loss of interaction or function at the protein level.

While the consequences of this mutation appear to not be as wide-ranging as the likely function of the *cut* gene, this mutation does have multiple effects on these bees. Both color and setal properties (length and thickness) are impacted by this phenotype and, while the whole body is not yellow, the effects occur in parts of the head, thorax, and upper abdomen. Furthermore, *cut* is known to play roles in reproductive success (e.g. sterility, semi-fertility) in both males and females of *Drosophila*, and to have effects on neurobiology. These functions may also be altered in the yellow mutant forms, as we have observed reduced reproductive success in the mutant, with queens exhibiting a worker-like behavior of being less inclined to mate and produce female progeny. This suggests some pleiotropy of this mutation, but also highlights a potential role of *cut* in driving caste-specific behaviors. A link between *cut* and caste specificity has been found in honeybees, where *cut*-like transcripts are downregulated in functionally sterile worker bees (which have only a few ovarioles) compared to its fertile and ovariole-rich queen counterparts [80]. Considering the negative fitness consequences for reproductive success, this particular protein-coding mutation would be selected against in nature.

Sanger sequencing of sister species revealed the implicated allele to be novel to this *B. terrestris* lab mutant, as it does not occur in similar phenotypes of closely related *Bombus s*.*s*. and is not known to occur in any Hymenopteran species. While our SNP is not found in these sister lineages and thus is not a result of sorting of ancestral recessive alleles, it is possible that other protein-coding region(s) or, more likely, cis-regulatory variants of *cut* occur in sister lineages to drive these phenotypes.

Experimental design of our research involves an approach utilizing genome-wide reduce-representation based sampling (RAD-Seq) followed by low-coverage whole-genome sequencing (WGS) of subsets of RAD-Seq samples (Figure 2d). RAD-Seq can work well to identify the chromosomal location of candidate locus (i.e. a distinct association peak(s) in GWAS) (e.g., [81], [82]), especially when recombination levels are low (e.g., in crosses), however, the subsampling of RAD-Seq most likely will miss specific functional SNPs. In our approach, a large number of samples are utilized in the first step (RAD-Seq) which, given the large linkage blocks present in crosses, clearly identified the chromosomal location of the candidate locus. Subsequent application of whole-genome sequencing of selected individuals representing the haplotype diversity apparent with RAD-Seq data across the parentage enabled us to localize the color locus down to the functional SNP, allowing success with a moderate amount of sampling (e.g., [82]) compared to large-scale GWAS in humans and other model organisms [83]. This works when the variant involves just a single allele, and thus applies to the search for a single fixed locus. color-related traits often are inherited in Mendelian fashion ([12], [82]) involving a single or a small number of large-effect loci [6]. Similar approaches involving both RAD-Seq and whole-genome sequencing were utilized in identifying the genetic basis of color polymorphism in butterflies [84], ladybird beetles [81], songbirds [82] and white tigers [73]. In addition to deploying a combination of RAD-Seq and WGS technique, we have also exploited the haplodiploid nature of sex determination of Hymenopterans in our experimental design as we used the drones (males) exclusively for genome sequencing approaches. Drones (males) are usually haploid thus heterozygosity and dominance effects on phenotypes are not possible and phasing is not required. Furthermore, sequence data from haploid samples are capable of producing good-quality *de novo* assemblies [85], genome alignment and variant calling datasets, even at moderate coverage (e.g., [12]). These properties make haploid systems particularly fruitful for genome-level discovery. These advantages enable lower-level sampling, especially valuable for under-researched Hymenoptera species where specimen collection can be difficult ([86], [87]), and considering the lack of substantial funding in non-model organisms in general [88].

## Conclusions

Through performing laboratory crosses in an emerging model bee, and utilizing a combination of large sample size RAD-Seq and low sample size whole genome sequencing, we have identified a new locus involved in insect color patterning that could also be involved in caste-specific fertility in bumble bees. This novel mutation reveals that protein-coding mutations in major developmental genes can have locally restricted effects. Future genomic research on natural yellow or black color variants in the *Bombus s*.*s*. lineage is needed to reveal whether this gene may be implicated in this color phenotype in nature, even if this particular mutation is not involved. Increasingly employed genome editing in Hymenoptera (e.g., [89], [90], [91]) as well as differential gene expression studies could be applied to better understand the role of *cut* and this particular mutation in bumble bees.

## Methods

### Sampled specimens and phenotypes

Initially a male with a yellow mutant color form containing yellow in the third thoracic segment, first abdominal segment, and posterior pleuron was generated from wildtype parents. Offspring from this individual were then crossed with wildtype *B. terrestris dalmatinus* and inbred across ∼18 generations to make a yellow mutant line. The yellow trait is recessive, with heterozygous females having wildtype coloration and her male offspring exhibiting both color forms. Yellow females showed reduced interest in mating with males of any phenotype or line (see Results), which resulted in reduced reproductive output of females and reproductive gynes. A few were produced, however, to maintain the line prior to experimentation, but not enough to maintain it for a longer term, as the line no longer exists. Selected specimens were examined for morphological differences across the body (e.g., wings, facial features, genitalia), including documentation of details of color pattern and setal morphology. Specimens were not examined internally.

Sixty wildtype *B. terrestris* queens from three colonies were placed in an arena and crossed with multiple (∼70) males from three colonies of the inbred yellow mutant line *B. terrestris* in a commercial bumble bee rearing facility (BizBee, EinYahav, Israel). Ten queens successfully generated offspring, from which eighty hybrid workers (F1) were generated. These workers were divided into four mini-colony worker groups (workers will lay male eggs when together in mini-groups, although not all individuals will lay) and allowed to generate male offspring (F2). Resulting F2 males exhibiting both wild-type and mutant color forms were selected for subsequent DNA sequencing. Given the pooled nature of the design, parentage could not be assessed, but should represent most of the ten original crosses.

### DNA extraction and Sequencing for RAD-Seq

Thoracic tissue from 90 haploid males (including 44 wildtype and 46 yellow mutant individuals), were extracted using a Qiagen DNeasy Blood and Tissue Kit with RNaseA treatment. Samples were extracted into a 250ul extraction buffer, dried using a SpeedVac, and re-eluted in 50ul water. Extracted samples were quantified using the Qubit BR dsRNA kit to ensure at least 250ng of DNA, and were assessed on a 1% agarose gel to ensure high-quality genomic DNA. Samples were prepared for RAD-Seq using protocols following [92] and [93]. This involved digesting 250ng of DNA per sample in a 50ul reaction with PstI-HF (NEB) for 1 hour at 37C, followed by 20 min. at 80C. Samples were divided into six sets (libraries) of 16, in each of which 16 different P1 barcoded adaptors were ligated using T4 DNA ligase with incubation for 22C for 1 hour, 65C for 10 min., and 30 min at room temperature. Each library was pooled and sheared to 300-700bp using a Covaris S2 sonicator. Gel-based size selection of 300-700bp fragments was then performed and samples purified using a Nucleospin Gel and PCR clean-up kit. Samples were end-repaired with the Quick Blunting Kit (NEB) and purified with Agencourt AMPure XP (Beckman Coulter) magnetic beads. This was followed by addition of dATP overhangs added with the Klenow exo (NEB), purification with AMpure XP beads, ligation of P2 adaptors, and additional purification with AMpure XP beads. Samples were then PCR amplified (98C for 30 sec, 16 cycles of 98C for 10sec and 72C for 1 min, 72C for 5 min) using Phusion High-Fidelity Master Mix with specific P2 Sanger indexing primers for each of the six pools. This process enabled the unique barcoding of all 9 samples. RAD-tag short read (2*150 bp, paired-end) libraries were sequenced using an Illumina HiSeq 2500 in Pennsylvania State University Genomics Core Facility (University Park, Pennsylvania, USA).

### DNA extraction and Sequencing for Whole-Genome Analysis

We selected a subset of samples for the whole-genome resequencing approach {both wildtype (n=7) and yellow mutant (n=7)} from the specimens used for RAD-Seq. This was performed given that RAD-Seq data revealed a peak of association but no fixed sites, thus functional SNPs were missed with this subsampling approach. To optimize the potential to find fixed sites within the association peak, we identified unique RAD-Seq haplotypes within the peak of association and selected individuals which maximally represented the range of differences in RAD-Seq haplotypes (Figure 2d). Genomic DNA extraction was conducted from thoracic tissues of haploid males using either Qiagen DNeasy or EZNA (Omega Bio-tek) kit followed up by RNaseA treatment. 150 bp paired-end sequencing libraries were prepared using Illumina TruSeq DNA Nano kits for whole-genome re-sequencing approach implemented in Illumina HiSeq 2500 sequencer (5-10 individuals per lane aimed at ∼30X per sample coverage) in Pennsylvania State University Genomics Core Facility (University Park, Pennsylvania, USA).

### Analysis of RAD-Seq Dataset

RAD-Seq analysis was completed using Stacks 2 software [94] following the recommendations of a published RAD-Seq analysis protocol [95]. First, the raw paired-end sequencing reads for six paired-end libraries were filtered and demultiplexed to generate individual-sample level sequence dataset using process_radtags unit of Stacks 2 software. Reads from individual samples (n=90, 44 wildtype and 46 yellow individuals) were aligned to the published *B. terrestris* genome assembly (Bter_1.0, GCA_000214255.1; [31]) using BWA aligner v. 0.7.17 [96] and the bwa-mem [97] algorithm with default parameters. After that, we ran pstacks, “cstacks”, “sstacks” and “gstacks” units of Stacks 2 as recommended by [95] and after applying a stringent filtering criteria (-p 2 -r 0.75; SNP must be present in both yellow and wildtype populations and genotyped in at least 75% of samples) generated a final SNP dataset of 57,712 SNPs using the “populations” unit of Stacks 2 that was ultimately used in genotype-phenotype association analysis. To test the association between case-control phenotype (wildtype and yellow mutants) with filtered SNP dataset, we performed a genotype-phenotype association testing utilizing Fisher’s exact test implemented in PLINK v. 1.9 [98]. Raw RAD-Seq reads for individual samples are available under NCBI BioProject [to be provided upon publication].

### Analysis of whole-genome resequencing dataset

Whole-genome resequencing data of wildtype (n=7) and yellow mutant (n=7) samples were analyzed using a previously implemented bioinformatic pipeline [12]. In brief, we applied appropriate adapter trimming (ILLUMINACLIP:adapters.fa:2:30:5), removal of low-quality bases (SLIDINGWINDOW:4:30 LEADING:3 TRAILING:3) and short-length (MINLEN:36) sequences using Trimmomatic v. 0.38 [99]. The published *B. terrestris* genome assembly (Bter_1.0, GCA_000214255.1; [31]) was used as a reference to align the trimmed reads using BWA aligner v. 0.7.17 [96] in bwa-mem [97] mode. Post-processing of aligned reads was implemented in SAMtools v. 1.8 [100] and Picard tools v. 1.119 [101]. We ran GATK v. 3.6 [102] in UnifiedGenotyper mode for multi-sample (n=14) variant calling using a haploidy-specific parameter (-ploidy 1 -glm SNP -stand_call_conf 25.0). Variant quality filtering was conducted in VCFtools v. 0.1.15 [103] using specific parameters (--max-missing 0.75 --minDP 3 --minQ 30 --min-alleles 2 --max-alleles 2), retaining high-quality biallelic SNPs and allowing no more than 25% missing data for any SNP position. The final genome-wide SNP dataset included 14,70,101 SNPs. Utilizing this filtered SNP dataset, we implemented Fisher’s exact test in PLINK 1.9 [98] to run a case-control (wildtype vs. yellow mutant phenotype) genotype-phenotype association analysis. Summary statistics of analyzed genomic samples are available in Supplementary Table 3 and raw genome sequencing reads are available under NCBI BioProject accession [to be provided upon publication].

### SNP Validation using Sanger Sequencing

To genotype the identified fixed SNP from the genotype-phenotype association analysis (see results) across specimens used in RAD-Seq analysis, genomic DNA was extracted from 83 individuals (42 wildtype and 41 yellow mutants) using thoracic tissue and standard protocols of a Qiagen DNeasy Kit. Primers were designed to amplify the candidate SNP-harboring region (SNP position 3802598 on NC_015770.1 identified from RAD-Seq and WGS analysis; Bter_CM1177_3802598_L 5’-CCTCTTTGTCCTTCGCTTGC-3’, Bter_CM1177_3802598_R 5’-CCAGCAAGATTCGCGAAATAGT-3’) and were amplified through PCR (94C 2 min, 35 cycles of [94C 30 sec, 51C 30 sec, 72C 90 sec], 72C 10 min; 15 ul reaction with 0.3ul 10uM primers, 5.9 ul water, 1 ul DNA, and 7.5ul HotStart Taq Mastermix (NEB)), purified with ExoSap-IT or a Qiagen MinElute PCR Purification Kit, and Sanger sequenced at the Pennsylvania State University Genomics Core Facility (University Park, Pennsylvania, USA). Manual inspection, trimming, and alignment of Sanger sequence data (chromatograms) was conducted in Geneious v. 8.1.9 [104] and the previously identified SNP was manually called.

### SNP comparison to other bumble bees and Hymenoptera

To add additional data toward understanding the fixation of the identified SNP by phenotype (SNP position 3802598 on NC_015770.1), we downloaded whole-genome sequencing data from 22 wildtype *B. terrestris* individuals available on NCBI (BioProject accession ID: PRJNA326162, sample names from I-D1 to I-D22) from [32]. Read trimming, alignment, alignment post-processing, multi-sample (n=22) variant calling, and variant filtering followed the bioinformatic pipeline for genomic data used above. The resulting SNP dataset (n=10,79,814) was visualized in IGV v. 2.3.86 [105] to identify whether the SNP remained fixed in the color locus considering their wildtype phenotype.

To test whether the identified color locus is implicated in wild populations of *B. terrestris* relatives, we included seven additional members of closely related species belonging to subgenus *Bombus s*.*s*. that vary in whether the above-mentioned segments are yellow or black (*B. patagiatus* (Yellow, Worker), *B. lucorum* (Black, Worker), *B. hypocrita* (Yellow, Male), *B. terricola* (Mostly Black, Queen), *B. sporadicus* (Yellow, Worker), *B. cryptarum s*.*s*. (Black, Worker), and *B. cryptarum moderatus* (Black, Worker); Figure 1d; Supplementary Table 1). Genomic DNA extraction, PCR, PCR purification, sequencing, sequence editing and allele calling followed protocols outlined above for genotyping of the narrowed locus.

To determine the nucleotide and amino-acid conservation of the implicated mutation more broadly across Hymenoptera, we performed multiple local-alignment searches (blastn, tblastn, blastx, blastp and tblastx) using NCBI BLAST [106] web interface [107] and downloaded gene orthologs (n=40) (Group 1911at7399) of all Hymenopterans from OrthoDB catalogue release 10.1 [46]. Amino acid sequences of orthologous proteins were aligned in Geneious v. 8.1.9 [104] using MAFFT alignment program [108] applying E-INS-i algorithm (default parameters: Scoring matrix: BLOSUM62, Gap Open Penalty 1.53, offset value 0) and the amino acid sequence conservation was visually compared.

### Gene annotation

To determine whether the SNP was located in a protein or cis-regulatory region, the SNP was compared against gene annotations available for *B. terrestris* on Hymenoptera Genome Database [109] where annotations have been made using automated approaches facilitated by transcriptome data, and the implicated region was further investigated through BLAST searches [106] to NCBI database to manually check this annotation. Although this supported the implicated SNP belonging to a protein-coding region, it suggested that the SNP was just three bases away from an intron-exon boundary. To check whether the SNP was contained in the final transcript and assess the potential for alternative splicing, we sequenced transcripts including the implicated region extracted from the head and abdominal tissues from one wildtype and one yellow mutant form. RNA extraction and DNA removal were conducted with the Direct-zol RNA Miniprep Plus kit following homogenization of tissue in 400ul Trizol using 4 metal beads in a 2ml tube in an Omni Bead Ruptor for 35 seconds. cDNA synthesis was conducted using a 15ul reduced volume reaction (includes 10.7 ul of 500ng RNA) but otherwise performed using standard protocols for the High Capacity cDNA Reverse Transcription Kit. Designed primers amplified a 230 bp fragment that spanned across three exons and two introns, including the intron nearest the implicated mutation (intron 1) (HHCutF_Exon1 5’ ACATTCAGGCCATGCAGTC 3’, HHCutR_Exon3 5’ GCTCTGCTGTAACCTGGACA 3’). PCR amplification was conducted using NEB 2X Hot Start Mastermix in a 15 ul reaction with the following conditions: 95C for 2 min, 30 cycles of 95C for 30 sec, 52C for 30 sec, 72C for 1 min, 5 min at 72C. A long-run high-density gel electrophoresis (2% agarose gel) was performed which confirmed just a single band for each PCR. Sanger sequencing of the PCR products (with HHCutF_Exon1 primer only) were conducted at the Pennsylvania State University Genomics Core Facility (University Park, Pennsylvania, USA). We utilized Geneious v. 8.1.9 [104] for manual inspection, trimming and alignment of Sanger sequence data (chromatograms) against publicly available *B. terrestris cut* cDNA sequences (NCBI Reference Sequence: XM_012311834.2).

### Protein structure and domain analysis

We tested whether the identified mutation alters structural and functional properties of the protein by performing translation analysis, predicting protein-protein interactions and structural domains of the identified protein, and determining whether predicted secondary structure is altered. Protein domains from Pfam [110] were assessed using default NCBI conserved domain database search [111], and coiled coils were predicted using the program MARCOIL with a threshold of 90 [112] from the HHPRED server toolkit [113]. Because the identified mutation (Ala38Pro) falls within the first predicted coiled coil region (amino acid residue range 30-63) of the *cut* protein, we repeated MARCOIL prediction using the SNP variant using the N-terminal 80 residues of each sequence. Potential protein interactions for *cut* were investigated using BIOGRID [51] for *Drosophila melanogaster* orthologs. *D. melanogaster* protein sequences reported to interact with *cut* were submitted to MARCOIL prediction, and sequence ranges of coiled-coils with probability above 90 were reported. *D. melanogaster* ortholog protein localization was reported from UniProtKB annotations [114]. PubMed identifiers from BIOGRID and *D. melanogaster* ortholog proteins from *B. terrestris* are reported for nuclear proteins with predicted coils that could potentially interact with the N-terminal coil in *cut*. We made structure models of the wildtype and yellow mutant coiled coil sequence (residues 20-55) using a helix identified from top template structure 6RW9_A (residues 2143-2178) with HHPRED [113].

## Declarations

### Ethics approval and consent to participate

Not applicable

### Consent for publication

Not applicable

## Availability of data and materials

Raw sequence reads for specimens used in RAD-Seq and WGS approaches are deposited under NCBI BioProject [to be provided upon publication]. cDNA sequences of *cut* sequences collected from head and abdominal tissues from wildtype and yellow mutant for testing alternative splicing in *B. terrestris* and cDNA sequences obtained for genotyping the color locus in *Bombus s*.*s*. are deposited in NCBI GenBank (accession numbers: [to be provided upon publication]). Datasets will be provided upon publication in the Dryad digital repository including final SNP datasets from publicly available wildtype *B. terrestris* sequencing data and in-house RAD-Seq and WGS based GWAS approaches along with associated population (phenotype) assignment files, MAFFT alignment file (nexus format) from protein homologs from OrthoDB database, manually edited alignment files (nexus format) from SNP validation in *B. terrestris* RAD-Seq samples.

## Authors’ contributions

HMH conceptualized the study, managed/conducted molecular extraction and analysis, performed phenotypic assays, performed some bioinformatic analysis, and supervised the project. SRR conducted the majority of bioinformatic and statistical analyses for WGS, RAD-Seq and comparative genomic approaches. LNK and NVG led protein structure analyses. JC discovered the mutant line, conducted crossing experiments, recorded behavioral observations, maintained bee colonies and selected bee samples. SRR and HMH curated the datasets. SRR wrote the manuscript, with contributions from all authors. All authors have contributed, read and approved the final manuscript.

## Acknowledgements

Thanks to Pennsylvania State University (PSU) Genomics Core Facility for sequencing services, and the PSU Institute for Computational and Data Sciences Advanced CyberInfrastructure (ICDS-ACI), where computational analysis was performed. Thanks to Benjamin Wadsworth, Rebecca Sommer, Tracy Baumgarten, Nick MacKnight, Briana Ezray, and Claudia Rosales for assistance in molecular techniques. Thanks to Sydney Cameron for providing sister taxa for sequencing. Thanks to Kaustubh Adhikari for assistance in GWAS plotting.

## Supplemental Information

**Supplementary Table 1:** Sample information for closely related species of *Bombus s*.*s*.

| Molecular ID | *Bombus* sp. | Caste/Sex | Color T3/M1 | Locality | Note<sub>1</sub> |
| --- | --- | --- | --- | --- | --- |
| BCP001 | *patagiatus* | Worker | Yellow | China: Sichuan, Tibetan Plateau, 3533m, 10-50km S. Hongyuan; 4-VIII-2002 |  |
| BCP003 | *lucorum* | Worker | Black | Switzerland: St. Gotthard's Pass; 27-VII-1999 |  |
| BCP005 | *sporadicus* | Worker | Yellow | Sweden: Abisko; 25-VII-1999 | SC193 [116] |
| BCP006 | *cryptarum moderatus* | Worker | Black | USA: Alaska | CP016A [17] |
| BCP007 | *cryptarum armeniacus* | Worker | Black | Turkey: Erzingan, Yeniyol-Ahmitli, 2120m; 5-VIII-2002 |  |
| BCP009 | *hypocrita* | Male | Yellow | Russia: Primorskiy Krai, Muravjov-Amurskii Peninsula; 25-VIII-2002 | SC168 [116] |
| Btrc042 | *terricola* | Queen | Mostly Black | USA: Pennsylvania, State College; 24-IV-2014 |  |

<sub>1</sub>Specimens are the same as those used in other studies, includes ID and reference from those studies.

**Supplementary Table 2 :** BIOGRID interactions for *D*. *melanogaster cut* protein

| NCBI Gene ID | Gene name | BIOGRID<sub>1</sub> | MARCOIL<sub>2</sub> | Localization<sub>3</sub> | PubMed ID | *B. terrestris* ortholog ID |
| --- | --- | --- | --- | --- | --- | --- |
| CG8597 | *lark* | Low | No | Nucleus, Cytoplasm | 27662615 [117] | NA |
| CG18005 | *beag* | High | 186-227 | Nucleus, spliceosome | 25242320 [118] | XP_012175085 |
| CG5874 | *nelf-A* | High | 277-304 | Nucleus, Chromosome | 25242320 [118] | XP_003401362 |
| CG14514 | *brd8* | High | 112-173 | Nucleus | 25242320 [118] | XP_012172412 |
| CG5994 | *nelf-E* | High | 8-36 | Nucleus, Chromosome | 25242320 [118] | XP_003396851 |
| CG8264 | *Bx42* | High | 296-330 | Nucleus | 25242320 [118] | XP_003397263 |
| CG16932 | *eps-15* | High | 423-552 | Extracellular | NA | NA |
| CG9638 | *ada2b* | High | No | NA | NA | NA |
| CG18297 | *cdk2ap1* | High | No | NA | NA | NA |
| CG17252 | *bcl7-like* | High | No | NA | NA | NA |
| CG1520 | *wasp* | High | No | NA | NA | NA |
| CG6521 | *stam* | High | No | NA | NA | NA |
| CG11482 | *mlh1* | High | No | NA | NA | NA |

<sub>1</sub>BIOGRID throughput1049 <sub>2</sub>Coiled-coil regions with MARCOIL prediction probability threshold >901050 <sub>3</sub>UniProtKB localization annotation

**Supplementary Table 3:**
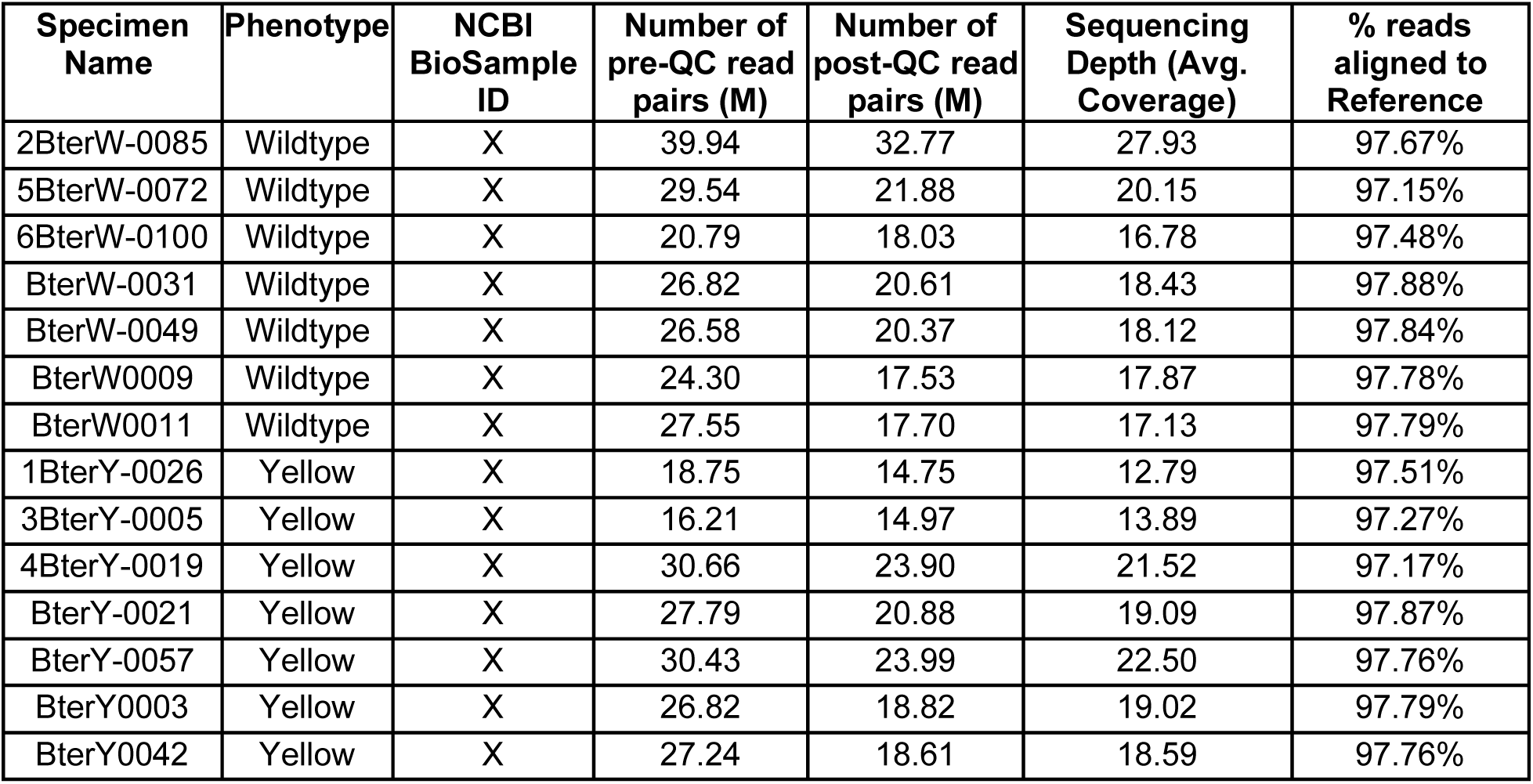
Summary Statistics for WGS Samples

## References

1. Letsou A, Bohmann D. Small flies—big discoveries: nearly a century of Drosophila genetics and development. Dev Dyn. Wiley Online Library; 2005;232:526–8.

2. Hales KG, Korey CA, Larracuente AM, Roberts DM. Genetics on the Fly: A Primer on the Drosophila Model System. Genetics. 2015;201:815–42.

3. Manceau M, Domingues VS, Linnen CR, Rosenblum EB, Hoekstra HE. Convergence in pigmentation at multiple levels: mutations, genes and function. Philos Trans R Soc Lond B Biol Sci. 2010;365:2439–50.

4. Yamamoto S, Jaiswal M, Charng W-L, Gambin T, Karaca E, Mirzaa G, et al. A drosophila genetic resource of mutants to study mechanisms underlying human genetic diseases. Cell. 2014;159:200–14.

5. St Johnston D. The art and design of genetic screens: Drosophila melanogaster. Nat Rev Genet. 2002;3:176–88.

6. Orteu A, Jiggins CD. The genomics of coloration provides insights into adaptive evolution. Nat Rev Genet. 2020;21:461–75.

7. Hines HM, Rahman SR. Evolutionary genetics in insect phenotypic radiations: the value of a comparative genomic approach. Curr Opin Insect Sci. 2019;36:90–5.

8. San-Jose LM, Roulin A. Genomics of coloration in natural animal populations. Philos Trans R Soc Lond B Biol Sci [Internet]. 2017;372. Available from: http://dx.doi.org/10.1098/rstb.2016.0337

9. Kronforst MR, Barsh GS, Kopp A, Mallet J, Monteiro A, Mullen SP, et al. Unraveling the thread of nature’s tapestry: the genetics of diversity and convergence in animal pigmentation [Internet]. Pigment Cell & Melanoma Research. 2012. p. 411–33. Available from: http://dx.doi.org/10.1111/j.1755-148x.2012.01014.x

10. Protas ME, Patel NH. Evolution of coloration patterns. Annu Rev Cell Dev Biol. 2008;24:425–46.

11. Reed RD, Papa R, Martin A, Hines HM, Counterman BA, Pardo-Diaz C, et al. optix drives the repeated convergent evolution of butterfly wing pattern mimicry. Science. 2011;333:1137–41.

12. Tian L, Rahman SR, Ezray BD, Franzini L, Strange JP, Lhomme P, et al. A homeotic shift late in development drives mimetic color variation in a bumble bee. Proc Natl Acad Sci U S A. 2019;116:11857–65.

13. Joron M, Frezal L, Jones RT, Chamberlain NL, Lee SF, Haag CR, et al. Chromosomal rearrangements maintain a polymorphic supergene controlling butterfly mimicry. Nature. 2011;477:203–6.

14. Nishikawa H, Iijima T, Kajitani R, Yamaguchi J, Ando T, Suzuki Y, et al. A genetic mechanism for female-limited Batesian mimicry in Papilio butterfly. Nat Genet. 2015;47:405–9.

15. Williams PH. An annotated checklist of bumble bees with an analysis of patterns of description (Hymenoptera: Apidae, Bombini). Bulletin-Natural History Museum Entomology Series. THE NATURAL HISTORY MUSEUM; 1998;67:79–152.

16. Williams P. The distribution of bumblebee colour patterns worldwide: possible significance for thermoregulation, crypsis, and warning mimicry. Biol J Linn Soc Lond. Oxford Academic; 2007;92:97–118.

17. Rapti Z, Duennes MA, Cameron SA. Defining the colour pattern phenotype in bumble bees (Bombus): a new model for evo devo. Biol J Linn Soc Lond. Oxford University Press; 2014;113:384–404.

18. Pimsler ML, Jackson JM, Lozier JD. Population genomics reveals a candidate gene involved in bumble bee pigmentation. Ecol Evol. 2017;7:3406–13.

19. Kraus FB, Szentgyörgyi H, Rozej E, Rhode M, Moroń D, Woyciechowski M, et al. Greenhouse bumblebees (Bombus terrestris) spread their genes into the wild. Conserv Genet. Springer; 2011;12:187–92.

20. Velthuis HHW, van Doorn A. A century of advances in bumblebee domestication and the economic and environmental aspects of its commercialization for pollination. Apidologie. EDP Sciences; 2006;37:421–51.

21. Kapheim KM, Pan H, Li C, Salzberg SL, Puiu D, Magoc T, et al. Genomic signatures of evolutionary transitions from solitary to group living. Science. American Association for the Advancement of Science; 2015;348:1139–43.

22. Harpur BA, Dey A, Albert JR, Patel S, Hines HM, Hasselmann M, et al. Queens and Workers Contribute Differently to Adaptive Evolution in Bumble Bees and Honey Bees. Genome Biol Evol. 2017;9:2395–402.

23. Li L, Su S, Perry CJ, Elphick MR, Chittka L, Søvik E. Large-scale transcriptome changes in the process of long-term visual memory formation in the bumblebee, Bombus terrestris. Sci Rep. 2018;8:534.

24. Stanley DA, Smith KE, Raine NE. Bumblebee learning and memory is impaired by chronic exposure to a neonicotinoid pesticide. Sci Rep. 2015;5:16508.

25. König C, Schmid-Hempel P. Foraging activity and immunocompetence in workers of the bumble bee, Bombus terrestris L. Proceedings of the Royal Society of London Series B: Biological Sciences. Royal Society; 1995;260:225–7.

26. Barribeau SM, Sadd BM, du Plessis L, Brown MJF, Buechel SD, Cappelle K, et al. A depauperate immune repertoire precedes evolution of sociality in bees. Genome Biol. 2015;16:83.

27. Matsumura C, Yokoyama J, Washitani I. Invasion status and potential ecological impacts of an invasive alien bumblebee, Bombus terrestris L.(Hymenoptera: Apidae) naturalized in Southern Hokkaido, Japan. GLOBAL ENVIRONMENTAL RESEARCH-ENGLISH EDITION-. 2004;8:51–66.

28. Knight ME, Martin AP, Bishop S, Osborne JL, Hale RJ, Sanderson RA, et al. An interspecific comparison of foraging range and nest density of four bumblebee (Bombus) species. Mol Ecol. 2005;14:1811–20.

29. Kraus FB, Wolf S, Moritz RFA. Male flight distance and population substructure in the bumblebee Bombus terrestris. J Anim Ecol. 2009;78:247–52.

30. Weidenmüller A. The control of nest climate in bumblebee (Bombus terrestris) colonies: interindividual variability and self reinforcement in fanning response. Behav Ecol. Oxford Academic; 2004;15:120–8.

31. Sadd BM, Barribeau SM, Bloch G, de Graaf DC, Dearden P, Elsik CG, et al. The genomes of two key bumblebee species with primitive eusocial organization. Genome Biol. 2015;16:76.

32. Liu H, Jia Y, Sun X, Tian D, Hurst LD, Yang S. Direct Determination of the Mutation Rate in the Bumblebee Reveals Evidence for Weak Recombination-Associated Mutation and an Approximate Rate Constancy in Insects [Internet]. Molecular Biology and Evolution. 2017. p. 119–30. Available from: http://dx.doi.org/10.1093/molbev/msw226

33. Harrison MC, Hammond RL, Mallon EB. Reproductive workers show queenlike gene expression in an intermediately eusocial insect, the buff-tailed bumble bee Bombus terrestris. Mol Ecol [Internet]. Wiley Online Library; 2015; Available from: https://onlinelibrary.wiley.com/doi/abs/10.1111/mec.13215?casa_token=a0XpctgwA88AAAAA:cbdHtwJP1XEoDRXbv1qiqITB85446gYI9e87rGAQ3ia3HIfd83qEeOIB28Poi8mnH_a4glcrvm7Uu1c

34. Zhao X, Xu W, Schaack S, Sun C. Genome-wide identification of accessible chromatin regions in bumblebee (Bombus terrestris) by ATAC-seq [Internet]. 2019 [cited 2020 Aug 7]. p. 818211. Available from: https://www.biorxiv.org/content/10.1101/818211v1.abstract

35. Williams PH, Brown MJF, Carolan JC, An J, Goulson D, Aytekin AM, et al. Unveiling cryptic species of the bumblebee subgenus Bombus s. str. worldwide with COI barcodes (Hymenoptera: Apidae). System Biodivers. Taylor & Francis; 2012;10:21–56.

36. Carolan JC, Murray TE, Fitzpatrick Ú, Crossley J, Schmidt H, Cederberg B, et al. Colour patterns do not diagnose species: quantitative evaluation of a DNA barcoded cryptic bumblebee complex. PLoS One. 2012;7:e29251.

37. Bossert S. Recognition and identification of bumblebee species in the Bombus lucorum-complex (Hymenoptera, Apidae) – A review and outlook. DEZ. Pensoft Publishers; 2015;62:19–28.

38. Rasmont P, Coppee A, Michez D, De Meulemeester T. An overview of the Bombus terrestris (L. 1758) subspecies (Hymenoptera: Apidae). Ann Soc Entomol Fr. Taylor & Francis; 2008;44:243–50.

39. Silva SE, Seabra SG, Carvalheiro LG, Nunes VL, Marabuto E, Mendes R, et al. Population genomics of Bombus terrestris reveals high but unstructured genetic diversity in a potential glacial refugium. Biol J Linn Soc Lond. Oxford Academic; 2019;129:259–72.

40. Hines HM, Witkowski P, Wilson JS, Wakamatsu K. Melanic variation underlies aposematic color variation in two hymenopteran mimicry systems. PLoS One. 2017;12:e0182135.

41. Polidori C, Jorge A, Ornosa C. Eumelanin and pheomelanin are predominant pigments in bumblebee (Apidae: Bombus) pubescence. PeerJ. 2017;5:e3300.

42. Hines HM. Bumble bees (Apidae: Bombus) through the ages: Historical biogeography and the evolution of color diversity. University of Illinois at Urbana-Champaign; 2008.

43. Massey JH, Wittkopp PJ. The Genetic Basis of Pigmentation Differences Within and Between Drosophila Species. Curr Top Dev Biol. 2016;119:27–61.

44. Kyrchanova O, Mogila V, Wolle D, Magbanua JP, White R, Georgiev P, et al. The boundary paradox in the Bithorax complex. Mech Dev. 2015;138 Pt 2:122–32.

45. Nijhout HF. Chapter 6 - Molecular and Physiological Basis of Colour Pattern Formation. In: Casas J, Simpson SJ, editors. Advances in Insect Physiology. Academic Press; 2010. p. 219–65.

46. Kriventseva EV, Kuznetsov D, Tegenfeldt F, Manni M, Dias R, Simão FA, et al. OrthoDB v10: sampling the diversity of animal, plant, fungal, protist, bacterial and viral genomes for evolutionary and functional annotations of orthologs [Internet]. Nucleic Acids Research. 2019. p. D807–11. Available from: http://dx.doi.org/10.1093/nar/gky1053

47. Pace CN, Scholtz JM. A helix propensity scale based on experimental studies of peptides and proteins. Biophys J. 1998;75:422–7.

48. O’Neil KT, DeGrado WF. A thermodynamic scale for the helix-forming tendencies of the commonly occurring amino acids. Science. 1990;250:646–51.

49. Surkont J, Pereira-Leal JB. Evolutionary patterns in coiled-coils. Genome Biol Evol. 2015;7:545–56.

50. Chang DK, Cheng SF, Trivedi VD, Lin KL. Proline affects oligomerization of a coiled coil by inducing a kink in a long helix. J Struct Biol. 1999;128:270–9.

51. Oughtred R, Stark C, Breitkreutz B-J, Rust J, Boucher L, Chang C, et al. The BioGRID interaction database: 2019 update. Nucleic Acids Res. 2019;47:D529–41.

52. Ebacher DJS, Todi SV, Eberl DF, Boekhoff-Falk GE. cut Mutant Drosophila Auditory Organs Differentiate Abnormally and Degenerate [Internet]. Fly. 2007. p. 86–94. Available from: http://dx.doi.org/10.4161/fly.4242

53. Blochlinger K, Bodmer R, Jack J, Jan LY, Jan YN. Primary structure and expression of a product from cut, a locus involved in specifying sensory organ identity in Drosophila. Nature. 1988;333:629–35.

54. Jack J, DeLotto Y. Structure and regulation of a complex locus: the cut gene of Drosophila. Genetics. 1995;139:1689–700.

55. Zhai Z, Ha N, Papagiannouli F, Hamacher-Brady A, Brady N, Sorge S, et al. Antagonistic regulation of apoptosis and differentiation by the Cut transcription factor represents a tumor-suppressing mechanism in Drosophila. PLoS Genet. 2012;8:e1002582.

56. Krupp JJ, Yaich LE, Wessells RJ, Bodmer R. Identification of genetic loci that interact with cut during Drosophila wing-margin development. Genetics. 2005;170:1775–95.

57. Eizirik E, Yuhki N, Johnson WE, Menotti-Raymond M, Hannah SS, O’Brien SJ. Molecular genetics and evolution of melanism in the cat family. Curr Biol. 2003;13:448–53.

58. Hoekstra HE, Hirschmann RJ, Bundey RA, Insel PA, Crossland JP. A single amino acid mutation contributes to adaptive beach mouse color pattern. Science. 2006;313:101–4.

59. Gratten J, Beraldi D, Lowder BV, McRae AF, Visscher PM, Pemberton JM, et al. Compelling evidence that a single nucleotide substitution in TYRP1 is responsible for coat-colour polymorphism in a free-living population of Soay sheep. Proc Biol Sci. 2007;274:619–26.

60. Werner T, Koshikawa S, Williams TM, Carroll SB. Generation of a novel wing colour pattern by the Wingless morphogen. Nature. 2010;464:1143–8.

61. Nijhout, F H. The development and evolution of butterfly wing patterns. Smithson Inst [Internet]. 1991 [cited 2020 Aug 7];293. Available from: https://ci.nii.ac.jp/naid/10003968117/

62. Sekimura T, Frederik Nijhout H. Diversity and Evolution of Butterfly Wing Patterns: An Integrative Approach. Springer; 2017.

63. Hines HM, Counterman BA, Papa R, Albuquerque de Moura P, Cardoso MZ, Linares M, et al. Wing patterning gene redefines the mimetic history of Heliconius butterflies. Proc Natl Acad Sci U S A. 2011;108:19666–71.

64. Martin A, Papa R, Nadeau NJ, Hill RI, Counterman BA, Halder G, et al. Diversification of complex butterfly wing patterns by repeated regulatory evolution of a Wnt ligand. Proc Natl Acad Sci U S A. 2012;109:12632–7.

65. Nadeau NJ, Pardo-Diaz C, Whibley A, Supple MA, Saenko SV, Wallbank RWR, et al. The gene cortex controls mimicry and crypsis in butterflies and moths. Nature. 2016;534:106–10.

66. Hannah AM. Radiation mutations involving the cut locus in Drosophila. Proceedings of the 8th International Congress of Genetics (Hereditas Suppl Vol) Stockholm. 1949. p. 588–9.

67. Lindsley DL, Grell EH. Genetic variations of Drosophila melanogaster. Publs Carnegie Instn; 1968.

68. Micchelli CA, Rulifson EJ, Blair SS. The function and regulation of cut expression on the wing margin of Drosophila: Notch, Wingless and a dominant negative role for Delta and Serrate. Development. 1997;124:1485–95.

69. Macdonald WP, Martin A, Reed RD. Butterfly wings shaped by a molecular cookie cutter: evolutionary radiation of lepidopteran wing shapes associated with a derived Cut/wingless wing margin boundary system [Internet]. Evolution & Development. 2010. p. 296–304. Available from: http://dx.doi.org/10.1111/j.1525-142x.2010.00415.x

70. Stern DL. Perspective: Evolutionary developmental biology and the problem of variation. Evolution. 2000;54:1079–91.

71. Carroll SB. Endless forms: the evolution of gene regulation and morphological diversity. Cell. 2000;101:577–80.

72. Van Belleghem SM, Rastas P, Papanicolaou A, Martin SH, Arias CF, Supple MA, et al. Complex modular architecture around a simple toolkit of wing pattern genes. Nat Ecol Evol. 2017;1:52.

73. Xu X, Dong G-X, Hu X-S, Miao L, Zhang X-L, Zhang D-L, et al. The genetic basis of white tigers. Curr Biol. 2013;23:1031–5.

74. Thurmond J, Goodman JL, Strelets VB, Attrill H, Gramates LS, Marygold SJ, et al. FlyBase 2.0: the next generation. Nucleic Acids Res. 2019;47:D759–65.

75. Bürglin TR, Affolter M. Homeodomain proteins: an update. Chromosoma. 2016;125:497–521.

76. Mier P, Alanis-Lobato G, Andrade-Navarro MA. Protein-protein interactions can be predicted using coiled coil co-evolution patterns. J Theor Biol. 2017;412:198–203.

77. Gillingham AK, Pfeifer AC, Munro S. CASP, the Alternatively Spliced Product of the Gene Encoding the CCAAT-Displacement Protein Transcription Factor, Is a Golgi Membrane Protein Related to Giantin [Internet]. Molecular Biology of the Cell. 2002. p. 3761–74. Available from: http://dx.doi.org/10.1091/mbc.e02-06-0349

78. Hoekstra HE, Coyne JA. The locus of evolution: evo devo and the genetics of adaptation. Evolution. 2007;61:995–1016.

79. Hsia CC, McGinnis W. Evolution of transcription factor function. Curr Opin Genet Dev. 2003;13:199–206.

80. Hepperle C, Hartfelder K. Differentially expressed regulatory genes in honey bee caste development. Naturwissenschaften. 2001;88:113–6.

81. Ando T, Matsuda T, Goto K, Hara K, Ito A, Hirata J, et al. Repeated inversions within a pannier intron drive diversification of intraspecific colour patterns of ladybird beetles. Nat Commun. 2018;9:3843.

82. Kim K-W, Jackson BC, Zhang H, Toews DPL, Taylor SA, Greig EI, et al. Genetics and evidence for balancing selection of a sex-linked colour polymorphism in a songbird. Nat Commun. 2019;10:1852.

83. Stranger BE, Stahl EA, Raj T. Progress and promise of genome-wide association studies for human complex trait genetics. Genetics. 2011;187:367–83.

84. Kunte K, Zhang W, Tenger-Trolander A, Palmer DH, Martin A, Reed RD, et al. doublesex is a mimicry supergene. Nature. 2014;507:229–32.

85. Iwasaki Y, Nishiki I, Nakamura Y, Yasuike M, Kai W, Nomura K, et al. Effective de novo assembly of fish genome using haploid larvae. Gene. 2016;576:644–9.

86. Sánchez-Bayo F, Wyckhuys KAG. Worldwide decline of the entomofauna: A review of its drivers. Biol Conserv. 2019;232:8–27.

87. Fischer B, Larson BMH. Collecting insects to conserve them: a call for ethical caution. Didham R, Gilbert F, editors. Insect Conserv Divers. 2019;12:173–82.

88. Sullivan W. The Institute for the Study of Non–Model Organisms and other fantasies. MBoC. American Society for Cell Biology (mboc); 2015;26:387–9.

89. Kohno H, Suenami S, Takeuchi H, Sasaki T, Kubo T. Production of Knockout Mutants by CRISPR/Cas9 in the European Honeybee, Apis mellifera L. Zoolog Sci. 2016;33:505–12.

90. Yan H, Opachaloemphan C, Mancini G, Yang H, Gallitto M, Mlejnek J, et al. An Engineered orco Mutation Produces Aberrant Social Behavior and Defective Neural Development in Ants. Cell. 2017;170:736-747.e9.

91. Li M, Au LYC, Douglah D, Chong A, White BJ, Ferree PM, et al. Generation of heritable germline mutations in the jewel wasp Nasonia vitripennis using CRISPR/Cas9. Sci Rep. 2017;7:901.

92. Heliconius Genome Consortium. Butterfly genome reveals promiscuous exchange of mimicry adaptations among species. Nature. 2012;487:94–8.

93. Baxter SW, Davey JW, Johnston JS, Shelton AM, Heckel DG, Jiggins CD, et al. Linkage mapping and comparative genomics using next-generation RAD sequencing of a non-model organism. PLoS One. 2011;6:e19315.

94. Catchen J, Hohenlohe PA, Bassham S, Amores A, Cresko WA. Stacks: an analysis tool set for population genomics. Mol Ecol. 2013;22:3124–40.

95. Rochette NC, Catchen JM. Deriving genotypes from RAD-seq short-read data using Stacks. Nat Protoc. 2017;12:2640–59.

96. Li H, Durbin R. Fast and accurate short read alignment with Burrows–Wheeler transform. Bioinformatics. Oxford Academic; 2009;25:1754–60.

97. Li H. Aligning sequence reads, clone sequences and assembly contigs with BWA-MEM [Internet]. arXiv [q-bio.GN]. 2013. Available from: http://arxiv.org/abs/1303.3997

98. Purcell S, Neale B, Todd-Brown K, Thomas L, Ferreira MAR, Bender D, et al. PLINK: a tool set for whole-genome association and population-based linkage analyses. Am J Hum Genet. 2007;81:559–75.

99. Bolger AM, Lohse M, Usadel B. Trimmomatic: a flexible trimmer for Illumina sequence data. Bioinformatics. 2014;30:2114–20.

100. Li H, Handsaker B, Wysoker A, Fennell T, Ruan J, Homer N, et al. The Sequence Alignment/Map format and SAMtools. Bioinformatics. 2009;25:2078–9.

101. Institute B. Picard tools. Broad Institute, GitHub repository; 2016.

102. McKenna A, Hanna M, Banks E, Sivachenko A, Cibulskis K, Kernytsky A, et al. The Genome Analysis Toolkit: a MapReduce framework for analyzing next-generation DNA sequencing data. Genome Res. 2010;20:1297–303.

103. Danecek P, Auton A, Abecasis G, Albers CA, Banks E, DePristo MA, et al. The variant call format and VCFtools. Bioinformatics. 2011;27:2156–8.

104. Kearse M, Moir R, Wilson A, Stones-Havas S, Cheung M, Sturrock S, et al. Geneious Basic: an integrated and extendable desktop software platform for the organization and analysis of sequence data. Bioinformatics. academic.oup.com; 2012;28:1647–9.

105. Robinson JT, Thorvaldsdóttir H, Winckler W, Guttman M, Lander ES, Getz G, et al. Integrative genomics viewer. Nat Biotechnol. 2011;29:24–6.

106. Altschul SF, Gish W, Miller W, Myers EW, Lipman DJ. Basic local alignment search tool. J Mol Biol. 1990;215:403–10.

107. Johnson M, Zaretskaya I, Raytselis Y, Merezhuk Y, McGinnis S, Madden TL. NCBI BLAST: a better web interface. Nucleic Acids Res. 2008;36:W5–9.

108. Katoh K, Standley DM. MAFFT multiple sequence alignment software version 7: improvements in performance and usability. Mol Biol Evol. 2013;30:772–80.

109. Munoz-Torres MC, Reese JT, Childers CP, Bennett AK, Sundaram JP, Childs KL, et al. Hymenoptera Genome Database: integrated community resources for insect species of the order Hymenoptera. Nucleic Acids Res. 2011;39:D658–62.

110. El-Gebali S, Mistry J, Bateman A, Eddy SR, Luciani A, Potter SC, et al. The Pfam protein families database in 2019. Nucleic Acids Res. 2019;47:D427–32.

111. Lu S, Wang J, Chitsaz F, Derbyshire MK, Geer RC, Gonzales NR, et al. CDD/SPARCLE: the conserved domain database in 2020. Nucleic Acids Res. 2020;48:D265–8.

112. Delorenzi M, Speed T. An HMM model for coiled-coil domains and a comparison with PSSM-based predictions. Bioinformatics. 2002;18:617–25.

113. Zimmermann L, Stephens A, Nam S-Z, Rau D, Kübler J, Lozajic M, et al. A Completely Reimplemented MPI Bioinformatics Toolkit with a New HHpred Server at its Core. J Mol Biol. 2018;430:2237–43.

114. UniProt Consortium. UniProt: a worldwide hub of protein knowledge. Nucleic Acids Res. 2019;47:D506–15.

115. gailhampshire. Bombus lucorum agg. male [Internet]. [cited 2020 Aug 7]. Available from: https://bit.ly/3kotdD5

116. Cameron SA, Hines HM, Williams PH. A comprehensive phylogeny of the bumble bees (Bombus) [Internet]. Biological Journal of the Linnean Society. 2007. p. 161–88. Available from: http://dx.doi.org/10.1111/j.1095-8312.2007.00784.x

117. Swain A, Misulovin Z, Pherson M, Gause M, Mihindukulasuriya K, Rickels RA, et al. Drosophila TDP-43 RNA-Binding Protein Facilitates Association of Sister Chromatid Cohesion Proteins with Genes, Enhancers and Polycomb Response Elements. PLoS Genet. 2016;12:e1006331.

118. Rhee DY, Cho D-Y, Zhai B, Slattery M, Ma L, Mintseris J, et al. Transcription factor networks in Drosophila melanogaster. Cell Rep. 2014;8:2031–43.

